# In Vitro Growth Effects of Morphine and Naloxone on Various Head and Neck Squamous Cell Cancer Cell Lines

**DOI:** 10.1101/320457

**Authors:** Stephen R Denton, Reema Padia, Jill E. Shea, Luke O Buchmann, Gregory J Stoddard, Matthew A Firpo, Tyler R Call

## Abstract

Understanding the potential effects of mu-agonism and antagonism on cancer cells is important for the perioperative physician. Previous studies suggest some tumor cells may have altered growth with mu-agonism or antagonism. This study investigates the effects of morphine (mu-agonist) and naloxone (mu-antagonist) in head and neck tumor cell lines (laryngeal squamous cell carcinoma (SCC), lateral tongue SCC and base of tongue SCC). Morphine showed no significant effect on tumor cell growth. Naloxone showed significant inhibition of growth in laryngeal SCC, but not in lateral or base of tongue SCC.

## Introduction

The perioperative period is a critical time for the recurrence and metastatic potential of cancer cells. (Abramovitch, Marikovsky, Meir, & Neeman, 1999; Ben-Eliyahu, 2003; Neeman & Ben-Eliyahu, 2013) Mu-agonists are routinely used in the perioperative period. Previous studies have suggested that the analgesic mu-agonist, morphine, may increase the growth of certain breast and lung cancer cells, possibly contributing to recurrence of these cancers. (Ecimovic et al., 2011; Mathew et al., 2011; Zylla, Kuskowski, Gupta, & Gupta, 2014) The aim of this study was to determine if exposing head and neck squamous cell cancer (SCC) cell lines to either a mu-agonist (morphine) or a mu-antagonist (naloxone) could have an effect on cell growth. These results could provide direction as to whether further investigation regarding the effects of mu-opioid agonists and antagonists on certain head and neck cancers might be warranted.

## Methods

SCC-12 laryngeal SCC, SCC-49 lateral oral tongue SCC, and SCC-74a base of tongue SCC cell lines were obtained from the University of Michigan SPORE bank laboratory and cultured in the University of Utah Surgical Laboratory. Resorufin reduction assays (alamarBlue) were used to monitor cell growth at 24, 48 and 72 hours when exposed to concentrations of morphine and naloxone separately, at 10, 25, 50, 75 and 100 ng/mL with a media-only control. Cells were grown using Dulbecco’s Modified Eagle’s Medium (cDMEM), 1 percent nonessential amino acids, 1 percent penicillin-streptomycin and 10 percent fetal bovine serum and kept in humidified atmosphere of 5 percent CO2 at 37 degrees Celsius. Ten passages were made to allow for sufficient cells. In order to measure cell growth and evaluate the effects of mu-opioid antagonism, 5x1000 cells/well in a ninety-six well culture plate were established. Six replicate wells were used per drug concentration with one column dedicated to the control with just media-exposed cells. The groups were: cells not exposed to any medications (control 1), and those exposed to 10 ng/mL, 25 ng/mL, 50 ng/mL, 75 ng/mL and 100 ng/mL of naloxone or morphine (Figure 1). Cell growth was evaluated by alamarBlue assay every 24 hours for 72 hours. The experiments were performed in triplicate. Concentrations of morphine and naloxone were selected base on the concentrations that may be seen in the standard clinical dosing.

**Figure 1.**
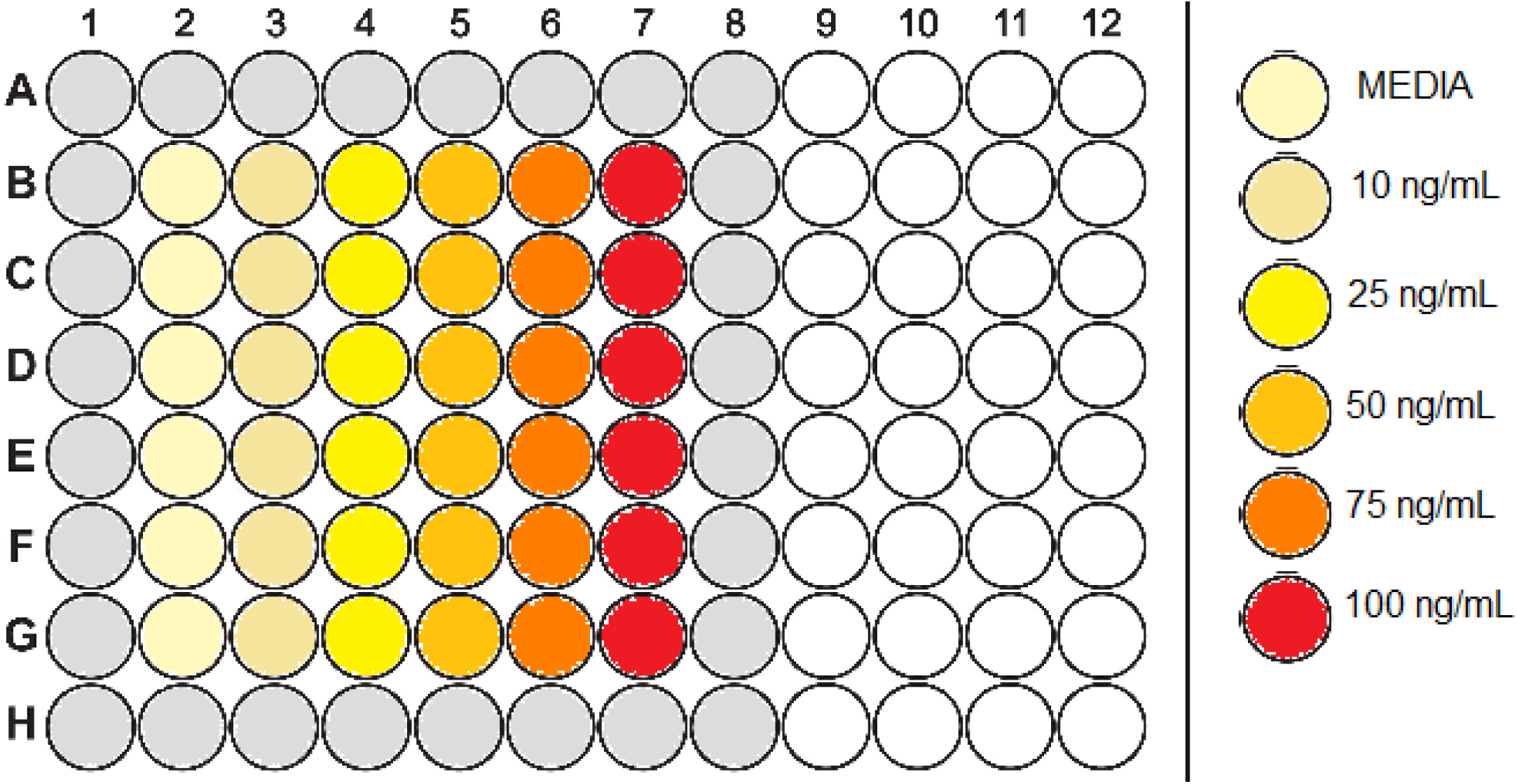
96-well plate layout for alamarBlue assay for naloxone and morphine.

## Results

Both the SCC-49 lateral oral tongue SCC and SCC-74 base of tongue SCC cell lines showed no changes in growth when exposed to either morphine or naloxone at the various concentrations. However, the SCC-12 laryngeal SCC showed a significant decrease in growth with exposure to naloxone at all concentrations except the 10ng/ml. (p < 0.001) The greatest inhibition was found at 48 hours (Figures 2 and 3). Morphine showed an increase in growth as shown in Figures 2 and 3, but the difference did not reach the level of significance.

**Figure 2.**
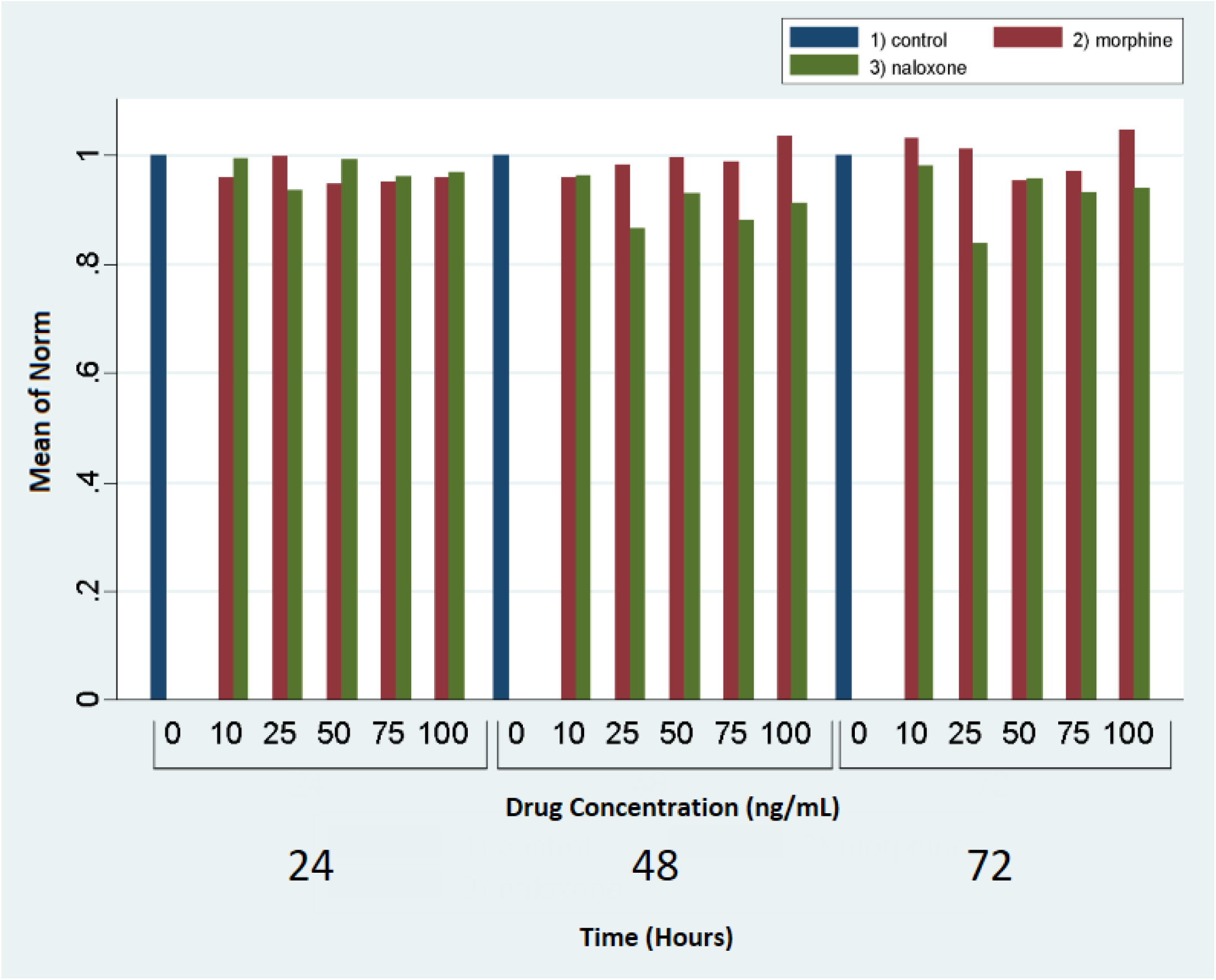
Comparison of growth of laryngeal SCC cells as compared to the control at 24, 48 and 72 hours when exposed to morphine and naloxone, separately.

**Figure 3.**
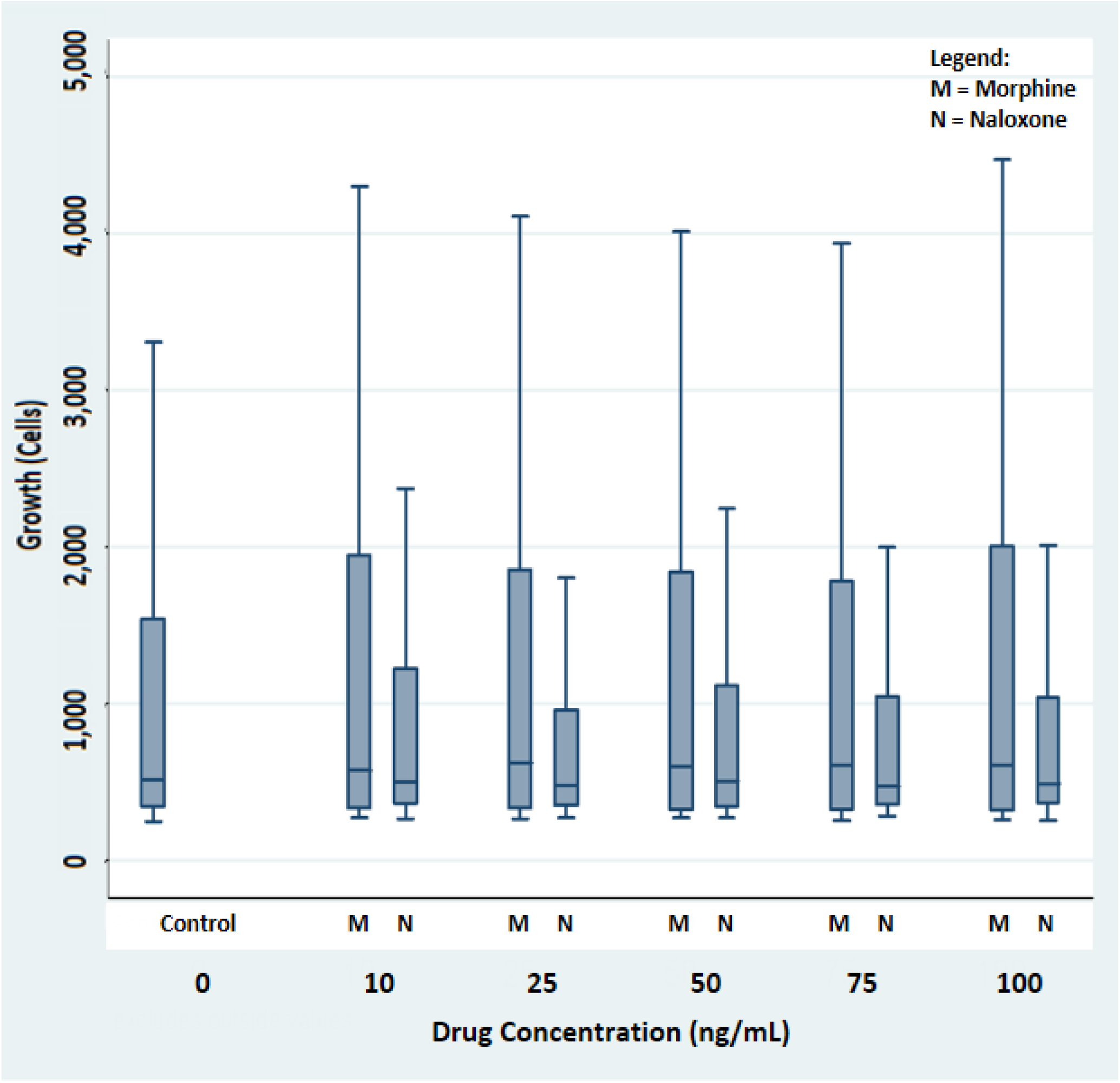
Comparison of morphine and naloxone exposed laryngeal SCC cell growth as compared to control values at 48 hours with increasing concentrations of the respective drug.

## Discussion

Opioid analgesics are commonly used in the head and neck cancer patient population and understanding the possible effects of mu-agonism and antagonism on these tumors is clinically important., The perioperative period is complex and many variables must be taken into account when providing care for the cancer patient. One of these variables is the direct effect of morphine (and other mu-agonists) on the tumor cells. These findings add a small but relevant piece of information to the larger perioperative picture. Although a trend in tumor cell growth was noted, statistical significance was not achieved and thus these findings do not support the idea that morphine directly promotes laryngeal or tongue SCC growth. The finding that mu-antagonism suppressed tumor cell growth while mu-agonism had no significant effect is peculiar. The mechanism by which laryngeal SCC tumor cell growth was repressed is unknown.

## Conclusion

No significant increase in tumor growth was noted in laryngeal SCC, base of tongue SCC or lateral tongue SCC when exposed to clinically relevant concentrations of morphine. Only laryngeal SCC showed an inhibitory growth effect when exposed to the mu-antagonist naloxone.

